# Frequency-dependent hybridization contributes to habitat segregation in monkeyflowers

**DOI:** 10.1101/2021.08.21.457231

**Authors:** Katherine Toll, David B. Lowry

## Abstract

Spatial segregation of closely related species is usually attributed to differences in stress tolerance and competitive ability. For both animals and plants, reproductive interactions between close relatives can impose a fitness cost that is more detrimental to the rarer species. Frequencydependent mating interactions may thus prevent the establishment of immigrants within heterospecific populations, maintaining spatial segregation of species. Despite strong spatial segregation in natural populations, two sympatric California monkeyflowers *(Mimulus nudatus* and *M. guttatus*) survive and reproduce in the other’s habitat when transplanted reciprocally. We hypothesized that a frequency-dependent mating disadvantage maintains spatial segregation of these monkeyflowers during natural immigration. To evaluate this hypothesis, we performed two field experiments. First, we experimentally added immigrants in varying numbers to sites dominated by heterospecifics. Second, we reciprocally transplanted arrays of varying resident and immigrant frequencies. Immigrant seed viability decreased with conspecific rarity for *M. guttatus*, but not *M. nudatus*. We observed immigrant minority disadvantage for both species, but driven by different factors– frequency-dependent hybridization for *M. guttatus*, and competition for resources and/or pollinators for *M. nudatus.* Overall, our results suggest a major role for reproductive interference in spatial segregation that should be evaluated along with stress tolerance and competitive ability.

## Introduction

Patterns of habitat segregation are common in closely related species and have been long mechanistically attributed to trade-offs between stress tolerance and competitive ability (Baker 1909, Connell 1961, Grace and Wetzel 1981, Yost et al. 2012, Peterson et al. 2013, Chen and Schemske 2015, Ferris and Willis 2018). Competition is well established to be an important mechanism driving zonation among closely related species with a high potential for niche overlap (Burns and Strauss 2011). However, mating between closely related species may also contribute to their spatial segregation (Anacker and Strauss 2014). The fitness costs of interspecific mating (reproductive interference) can drive patterns of segregation or coexistence through demographic displacement (Groning and Hochkirch 2008) or the evolution of increased pre-zygotic isolation in sympatry (Servedio and Noor 2003). The ecological and evolutionary outcomes of costly interspecific mating – spatial segregation (Groning and Hochkirch 2008) or character displacement (Brown and Wilson 1956) – depend on the relative rate of evolutionary change versus the rate of demographic decline in sympatry (Kyogoku and Wheatcroft 2020). While the initial establishment of habitat segregation could result from exclusion in sympatry or evolve directly in response to reproductive interference (Kyogoku and Kokko 2020), the longterm maintenance of zonation patterns requires mechanisms that result in the demographic decline of rare immigrants.

Plant species that co-occur and overlap in flowering often compete for pollinators and often suffer consequent fitness reductions (Mitchell et al. 2009). Competition for pollination can reduce fitness through reductions in visitation or through interspecific pollen transfer (Waser 1978a, b). Both mechanisms of competition for pollination can affect fitness in a frequencydependent manner. Rarer species may receive fewer visits and are expected to receive less conspecific pollen and more heterospecific pollen than common species. Models of reproductive interference thus predict fitness declines with increasing rarity (Levin and Anderson 1970, Kuno 1992). Rare-species disadvantage caused by reproductive interference is a major factor limiting persistence of polyploid species, which arise in populations of their diploid progenitors (minority cytotype exclusion, Levin 1975). Thus, considerable focus has been placed on these dynamics in co-occurring diploid and polyploid species (Husband 2000, Baack 2005, Buggs and Pannell 2006). However, the conditions experienced by rare polyploids are similar to that of parapatric species that meet in secondary contact (Lewis 1961, Ribeiro and Spielman 1986, Thum 2007), immigrants of species that are micro-spatially segregated within regions of sympatry (Singer 1990, Friberg et al 2013, Christie and Strauss 2020), or nonnative species interacting with native congeners (Takakura et al. 2009, Takakura and Fujii 2010, Takakura 2013, Takakura and Fujii 2015).

An extreme cost to reproductive interactions is the production of inviable seeds from interspecific pollen transfer. This hybrid seed inviability is a common reproductive isolating barrier in plants and particularly among species in the *Mimulus guttatus* species complex (Coughlan et al. 2020, Garner et al. 2016, Oneal et al. 2016, Sandstedt et al. 2021, Vickery 1978). Inviable hybrid seeds produced between species in the *M. guttatus* species complex have a characteristic flat or shriveled shape, which is caused by arrested endosperm development. These inviable seeds can be distinguished by eye from round viable seeds with fully formed endosperm produced by conspecific pollinations.

Populations of the widespread *M. guttatus* have repeatedly evolved tolerance to harsh serpentine soils (Selby and Willis 2018) and co-occur with the geographically restricted serpentine soil endemic *M. nudatus* in the Northern coast range of California. *M. guttatus* typically grows in streams, seeps, or meadows, whereas *M. nudatus* typically grows in washes, on rock outcroppings, or drier rocky areas adjacent to the streams inhabited by *M. guttatus. M. nudatus* typically grows in barer habitats with fewer co-occurring con- and heterospecifics than *M. guttatus*, a common pattern in serpentine soil endemics in contrast to species that occur both on and off serpentine soils (Sianta and Kay 2019). Since habitats occupied by *M. guttatus* and *M. nudatus* are often in close proximity (within meters) and immigrant individuals are regularly found, dispersal limitation does not explain spatial segregation in these species. When transplanted reciprocally at equal frequencies, *M. guttatus* has higher survival and produces more seed than *M. nudatus* in both habitats, demonstrating that stress tolerance or competition are not sufficient to explain patterns of spatial segregation between these species (Toll and Willis 2018). In those reciprocal transplants, both species produced many inviable F1 hybrid seeds as immigrants, demonstrating the occurrence of reproductive interference (Toll and Willis 2018). While a combination of factors limits the persistence of *M. nudatus* immigrants in habitats occupied by *M. guttatus*, including a lack of tolerance to flooding in *M. nudatus* (Toll and Willis 2018) and inbreeding depression (Toll et al. 2021), these factors do not sufficiently explain the absence of *M. guttatus* in the drier habitats occupied by *M. nudatus.* We therefore hypothesize that frequency-dependent reproductive interference may maintain this spatial segregation pattern by imposing a strong fitness cost on natural immigrants, which usually occur alone or in very small numbers.

Using observational and experimental data collected across two field seasons, we asked three questions in this study. First, does hybridization reduce fecundity when *M. guttatus* and *M. nudatus* naturally occur in close proximity? If hybridization contributes to the maintenance of spatial segregation, we expect that seed viability will increase with increasing conspecific frequency. Second, when species immigrate to sites occupied by heterospecifics, do components of fitness in immigrants or residents (flowers per plant, seeds per flower, and seed viability) increase with their relative frequency? If resource competition contributes to the maintenance of habitat segregation, we expect that flowers per plant will increase with increasing conspecific frequency. If a lack of conspecific mates or pollinator competition contributes to the maintenance of habitat segregation, we expect that seeds per flower will increase with increasing conspecific frequency. If hybridization contributes to the maintenance of habitat segregation, we expect that seed viability will increase with increasing conspecific frequency. Third, does lifetime fecundity (measured as viable seeds per plant) in immigrants or residents increase with increasing conspecific frequency in experimental sympatry? If minority disadvantage contributes to the maintenance of habitat segregation, we would expect that viable seeds per immigrant or resident plant to increase with their increasing frequency, changing the relative performance of immigrants relative to residents.

## Materials and methods

### Reproductive interference in natural populations: an observational transect

To characterize the existence and possible magnitude of reproductive interference in cooccurring populations of *M. guttatus* and *M. nudatus,* we surveyed plants along a transect spanning an exposed serpentine outcrop that was inhabited by a pure stand of *M. nudatus,* a transition zone where both species were present at lower density, terminating in a serpentine seep with a pure stand of *M. guttatus* (38°51.528’ N, 122°24.614’ W). The entire length of the transect, spanning the two pure stands through a small zone of overlap, was six meters. We counted all the fruit produced by each species within eight 42 × 66 cm gridded quadrats centered on the transect line, with the long side parallel to the transect line and evenly spaced along the length of the transect. We counted all fruit present within the quadrat at 7cm intervals along the long side of the quadrat grid. At approximately 21cm intervals along the long side of the quadrat, we collected one plant if only one species was present (n=13 *M. nudatus* and n=9 *M. guttatus)* or two plants if both species were present (n=10 *M. nudatus* and n=10 *M. guttatus)* at the mid-point of the short side of the quadrat. We counted fruit numbers instead of species abundances because flower number, not plant number, is what pollinators see in dense overlapping stands (Gardner and MacNair 2000). We counted all seeds produced by the plants collected at 21 cm intervals and categorized them as round, viable (conspecific) or flat, inviable F1 hybrid seeds based on morphology (Oneal et al. 2016). We calculated seed viability by dividing the number of round viable conspecific seeds by the total number of seeds produced by each plant. We estimated Pearson correlation coefficients for the relationship between the proportion of conspecific fruit and seed viability per sampled plant in R v 4.1.2 (R Core Team 2021).

### Experimental Transplants

To test whether frequency-dependence contributes to the maintenance of habitat segregation, we performed two reciprocal transplants over two years. In the first field season (2018), we transplanted immigrants in varying abundances into natural populations of resident heterospecifics. This allowed us to simulate a situation where immigrants arrive at low abundances to sites already established by heterospecifics. However, since we did not transplant the naturally occurring resident species with the immigrants, we lacked an appropriate comparison for immigrant performance relative to residents. In the second field season (2019), we transplanted immigrants and residents at varying frequencies reciprocally, allowing us to compare the overall fecundities of experimental immigrants and residents.

#### M. guttatus immigration experiment

In February 2018, we transplanted variable numbers of immigrants into naturally dense patches of native seedlings (>100 per half meter). While we performed this experiment reciprocally, all but nine immigrant *M. nudatus* transplants died before flowering. We lacked sufficient seedlings to replace them, and they were not analyzed further. Therefore, this experiment only allowed us to examine the immigration dynamics of *M. guttatus* into *M. nudatus* habitat (the direction less-well explained by our previous studies).

We collected seeds from a local population of *M. guttatus* at the University of California McLaughlin reserve (38°51.528’ N, 122°24.614’ W) in 2013. We grew seeds from field-collected maternal families in a greenhouse and crossed a single plant from each family to a single plant from a different family to produce outbred maternal families in 2013 (described in more detail in Toll et al. 2021). To minimize effects of inbreeding on fecundity (see Toll et al. 2021), we germinated 6 outbred maternal families of *M. guttatus* to use as focal plants. These plants were initially grown in the shade house at the UC McLaughlin reserve in 2018. We did not have sufficient seed to use these same outbred lines as neighbors, so we pooled equal numbers of seed from twenty field-collected maternal families of *M. guttatus* to germinate in the shade house to use as neighbors. We planted 72 blocks of focal *M. guttatus* plants in random order by maternal family, then randomly assigned each family to one of the 6 levels of the initial conspecific neighbor treatments (0, 8, 24, 48, 80, and 120 neighbors, Figure 1A). We transplanted a range of *M. guttatus* seedlings that were ecologically relevant if a single fruit dispersed into *M. nudatus’* habitat, given that each fruit produces an average of 200 seeds (Toll and Willis 2018). *M. guttatus* seedlings were planted within naturally dense patches of *M. nudatus* (>100 native *M. nudatus* within a half meter of each focal *M. guttatus* plant, Figure 1B). We planted *M. guttatus* neighbors within a half-meter of each focal plant in a regularly spaced array and we planted these blocks over three meters away from one another.

**Figure 1.**
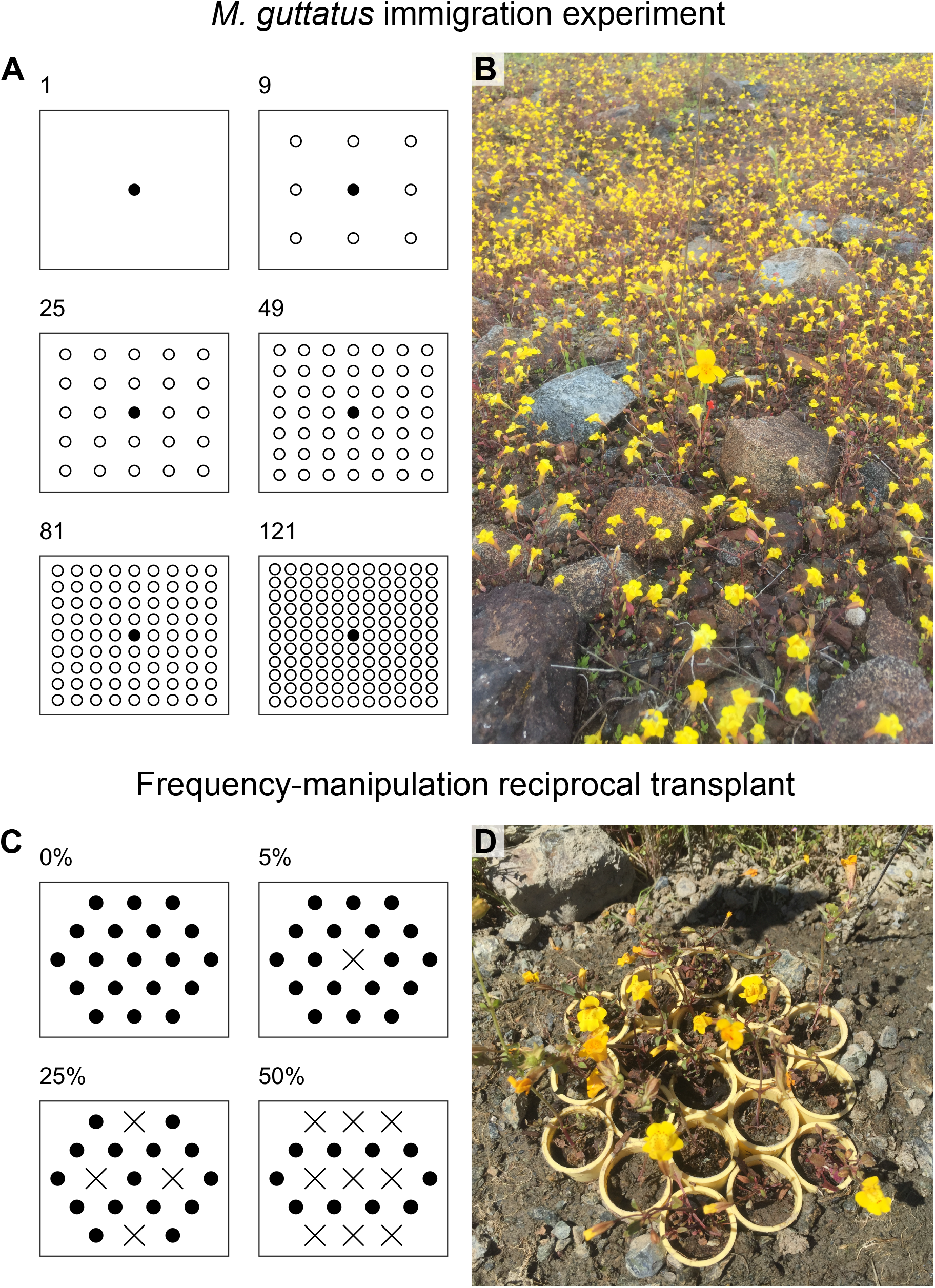
(A) Schematic of the 2018 *M. guttatus* immigration experiment showing positions of focal immigrant *M. guttatus* plants (filled circle) and conspecific neighbors (open circles) within each block. (B) Photograph of an *M. guttatus* transplant with no conspecific neighbors, surrounded by native *M. nudatus.* (C) Schematic of the 2019 frequency-manipulation reciprocal transplant experiment showing the showing positions of immigrant (X) and resident (o) plants within each immigrant frequency treatment. Figure adapted from Nagy (1997). Five replicates per treatment were planted at each site, resulting in 20 blocks containing residents and 15 blocks containing immigrants at each site. (D) Photograph of an example 50% immigrant (9N:10G) plot in the seep site. The diameter of each pot was 2.54 cm and the largest diameter of each array was 12.7cm. Native *Mimulus* plants within 0.25m of each array were removed prior to transplanting.

We replaced *M. guttatus* focal transplants as they died of transplant shock, resulting in 62 focal plants (of the original 72 plus 28 replacements) that survived to flower by the end of the season. Neighbor transplant mortality rates were also high; 78% of *M. guttatus* transplants did not survive to flowering, resulting in a range of flowering neighbors from 0 to 103, with a median of 1 neighbor surviving to flowering. In subsequent analyses of the effect of conspecific neighbors on components of fitness, we only included neighbors that successfully survived to flower, as only they could potentially reduce reproductive interference. Mortality rates from this transplant experiment are consistent with previous estimates at this site: 73%, 33%, and 82% of *M. guttatus* seedlings died at this site in 2015, 2016, and 2017 respectively (site “QV” in Toll and Willis 2018). Thus, we are confident that our transplants represent an ecologically realistic range of potential surviving immigrant *M. guttatus* seedlings into *M. nudatus* habitat. We counted all seeds produced by the 62 focal plants that produced fruit and categorized them as viable round conspecific-pollinated or inviable flat heterospecific-pollinated (hybrid) seeds based on morphology.

##### Statistical analysis

To examine how the number of conspecific neighbors affected focal plant fitness components (flowers per plant, total seed per flower, and seed viability per focal *M. guttatus* transplant) we used generalized linear mixed models in the R package *glmmTMB* (Brooks et al. 2017). We fit models with each fitness component as the dependent variable, the number of surviving conspecific neighbors as a fixed effect, and maternal family (n=6) of the focal plant as a random effect. Models of total seeds per flower had an additional nested random effect term for individual plant (n=62) since each focal plant produced multiple flowers.

We fit each count model first with a Poisson error distribution and identified the best fitting error distribution by evaluating model diagnostics with the R package DHARMa (Hartig 2019). The effect of conspecific neighbors on flowers per focal transplant was estimated using a negative binomial model because Poisson models were significantly overdispersed. The effect of conspecific neighbors on seeds per flower was estimated using zero-inflated negative binomial models with a constant intercept because Poisson and negative binomial models were significantly overdispersed and zero-inflated. The effect of conspecific neighbors on seed viability was estimated using a beta-binomial model because the binomial model was significantly overdispersed.

We assessed the significance of the fixed effect (the number of conspecific neighbors) with the best fitting model for each fitness component using Wald Type II Chi-squared tests (R package *car,* Fox and Weisberg 2019). We predicted marginal effects and bootstrapped (n=500 iterations) 95% confidence intervals with the R package *ggeffects* (Lüdecke 2018). We plotted raw data and predictions with *ggplot2* (Wickham 2016) and combined plots with *patchwork* (Pederson 2020).

#### Frequency-manipulation reciprocal transplant

To test whether frequency-dependence contributes to the maintenance of habitat segregation, we reciprocally transplanted *M. guttatus* and *M. nudatus* at different frequencies but with the same number of individuals in experimental patches. These mixed species patches were transplanted into a seep dominated by *M. guttatus* and a wash dominated by *M. nudatus*. The plants used in this experiment were all collected directly from natural populations at each transplant site in the year (2019) of the experiment. In April 2019, we dug up 380 seedlings per species from each site and repotted them in yellow 49-mL (2.54 cm diameter) Cone-tainers^TM^ filled with potting soil (Stuewe and Sons, Inc., Tangent, OR, USA). We maintained seedlings in a shadehouse at the UC McLaughlin Reserve for a week prior to redistributing seedlings in Cone-tainers^TM^ at both sites. We sunk Cone-tainers^TM^ into the ground, so the top was flush with the soil surface at each site in 19 plant hexagonal arrays (Figure 1C). Arrays were planted within natural populations of each species, but we cleared native *Mimulus* plants within 0.25 m of each array. We designed this experiment following the frequency manipulation arrays from Nagy (1997): 0% immigrants (0 immigrants, 19 residents); 5% immigrants (1 immigrant, 18 residents); ~25% immigrants (4 immigrants, 15 residents); ~50% immigrants (9 immigrants, 10 residents) (Figure 1D). Higher levels of immigrants than natives would have been unrealistic in this system. For clarity, we abbreviate frequency treatments as the ratio of *M. guttatus* (G) and *M. nudatus* (N) transplants. We assigned plants to four frequency treatments in the seep site (19G, 18G: 1N, 15G:4N, and 10G:9N) which is naturally occupied by *M. guttatus,* and four frequency treatments in the wash site (19N, 18N:1G, 15N:4G, and 10N:9G) which is naturally occupied by *M. nudatus.* We planted 5 replicates per frequency treatment per site, for a total of 20 blocks per site and 40 blocks overall. Blocks were planted at least three meters away from each other. We added supplemental water to blocks for a month after transplanting and replaced seedlings that died prior to flowering to ensure that the same number of plants flowered in each frequency treatment. We left containers in the field until all plants terminated flowering and ripened their fruit. We collected fruit from each transplant in the field prior to digging up containers. We counted and categorized all seeds produced by experimental plants and estimated the hybridization rate using the same methods as the observational transect. We estimated lifetime fecundity (viable seeds per plant) from three fitness components: the total number of flowers produced per plant, the total number of seeds produced per flower, and seed viability (=viable seeds/total seeds produced).

##### Statistical analysis

To examine how the immigrant frequency treatments affected plant fitness components (flowers per plant, total seed per flower, and seed viability) and lifetime fecundity (viable seeds per plant), we used generalized linear mixed models in the R package *glmmTMB* (Brooks et al. 2017). We tested whether fitness components differed among frequency treatments for each species in each site separately because no immigrants were transplanted in the 0% immigrant frequency treatment (Figure 1D). We fit models with each fitness component as the dependent variable, immigrant frequency as a fixed effect and block as a random effect (block n=15 for immigrants, n=20 for residents). We also tested whether lifetime fecundity differed between species, but only within the treatments where both immigrants and residents were present (5%, 25%, and 50% immigrant frequency treatments) in each site separately. We fit models with lifetime fecundity as the dependent variable, immigrant frequency, species and their interaction as fixed effects and block as a random effect (block n=15).

Poisson models of count fitness components and lifetime fecundity were significantly overdispersed, and models of seeds per flower and viable seeds per plant were significantly zero-inflated. Thus, the effect of immigrant frequency on flowers per plant was estimated using a negative binomial distribution, while total seeds per flower and viable seeds per plant were estimated using a zero-inflated negative binomial distribution with a constant zero-inflation intercept applied to all observations. The effect of immigrant frequency on seed viability was analyzed using a binomial model for immigrants, and a beta-binomial model for residents because binomial models for residents were significantly overdispersed.

We assessed the significance of the fixed effect (immigrant frequency treatment) with the best fitting model for each fitness component using Wald Type II Chi-squared tests and assessed the significance of the fixed effects (immigrant frequency treatment, species, and their interaction) for lifetime fecundity using Wald Type III Chi-squared tests (R package *car,* Fox and Weisberg 2019). We tested whether fitness components differed among treatments within species, and whether species differed in lifetime fecundity within each frequency treatment with Tukey post-hoc contrasts (R package *emmeans*, Lenth 2020). We estimated marginal means and bootstrapped (n=500 iterations) 95% confidence intervals with the R package *ggeffects* (Lüdecke 2018). We plotted marginal means with *ggplot2* (Wickham 2016) and combined plots with *patchwork* (Pederson 2020).

## Results

### Reproductive interference in natural populations: an observational transect

Individuals collected along a transect produced a high proportion of viable seeds in zones where only conspecifics were present (average 0.92 for *M. guttatus;* 0.84 for *M. nudatus),* even though heterospecifics were in very close proximity (always within 3m). In contrast, individuals collected in the 3m overlap zone of the two species produced far lower proportions of viable seeds (average 0.53 in *M. guttatus;* 0.66 in *M. nudatus,* Figure 2). Seed viability was positively correlated with the local frequency of conspecific fruit for *M. guttatus* plant sampled along the transect (Pearson correlation coefficient= 0.66; 95% CI= 0.29, 0.86; t = 3.61, df=17, *p* = 0.002), meaning that greater numbers of adjacent conspecific flowers and fruits were associated with greater proportions of viable seeds. However, seed viability was not significantly correlated with the local frequency of conspecific fruit for *M. nudatus* plants sampled along the transect (Pearson correlation coefficient= 0.18; 95% CI= −0.28, 0.57; t = 0.79; df = 19; *p* = 0.44). These data from a natural transition zone suggest that reproductive interference is more likely to be frequencydependent in *M. guttatus* immigrants than in *M. nudatus* immigrants.

**Figure 2.**
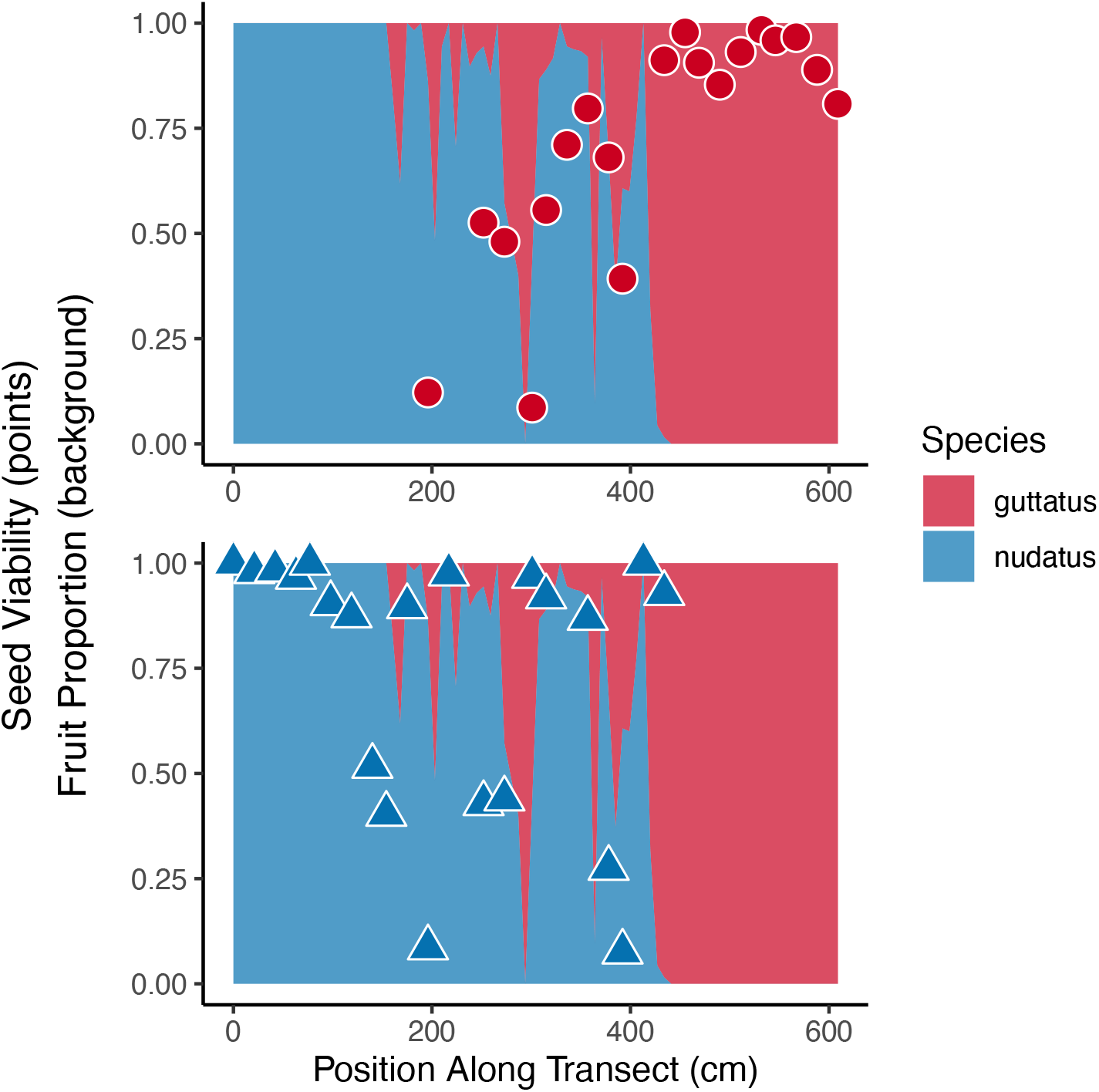
Transect of co-occurring *M. nudatus* and *M. guttatus.* Background colors depict the frequency of *M. nudatus* fruit (blue) vs *M. guttatus* fruit (red). Points indicate the proportion of inviable F1 hybrid seeds produced by individual *M. nudatus* (triangles) and *M. guttatus* (circles) plants collected every 21cm along the transect.

### Experimental Transplants

#### M. guttatus immigration experiment

As expected from the observational transect, *M. guttatus* immigrant transplants hybridized less and produced a greater proportion of viable conspecific seeds when surrounded by more flowering conspecific neighbor transplants (Figure 3C), as would occur if an intact fruit or multiple seeds colonized a habitat patch occupied by heterospecifics. Seed viability increased with increasing numbers of conspecific neighbors (Wald type II Chi-square test: χ2 = 7.49, df = 1, *p* = 0.006; coefficient estimate = 0.023, SE = 0.009, z-value = 2.74, *p* = 0.006). The model-predicted seed viability for rare immigrants increased from 0.24 for immigrants without conspecific neighbors, to 0.78 in the plant with 103 neighbors (Figure 3C).

**Figure 3.**
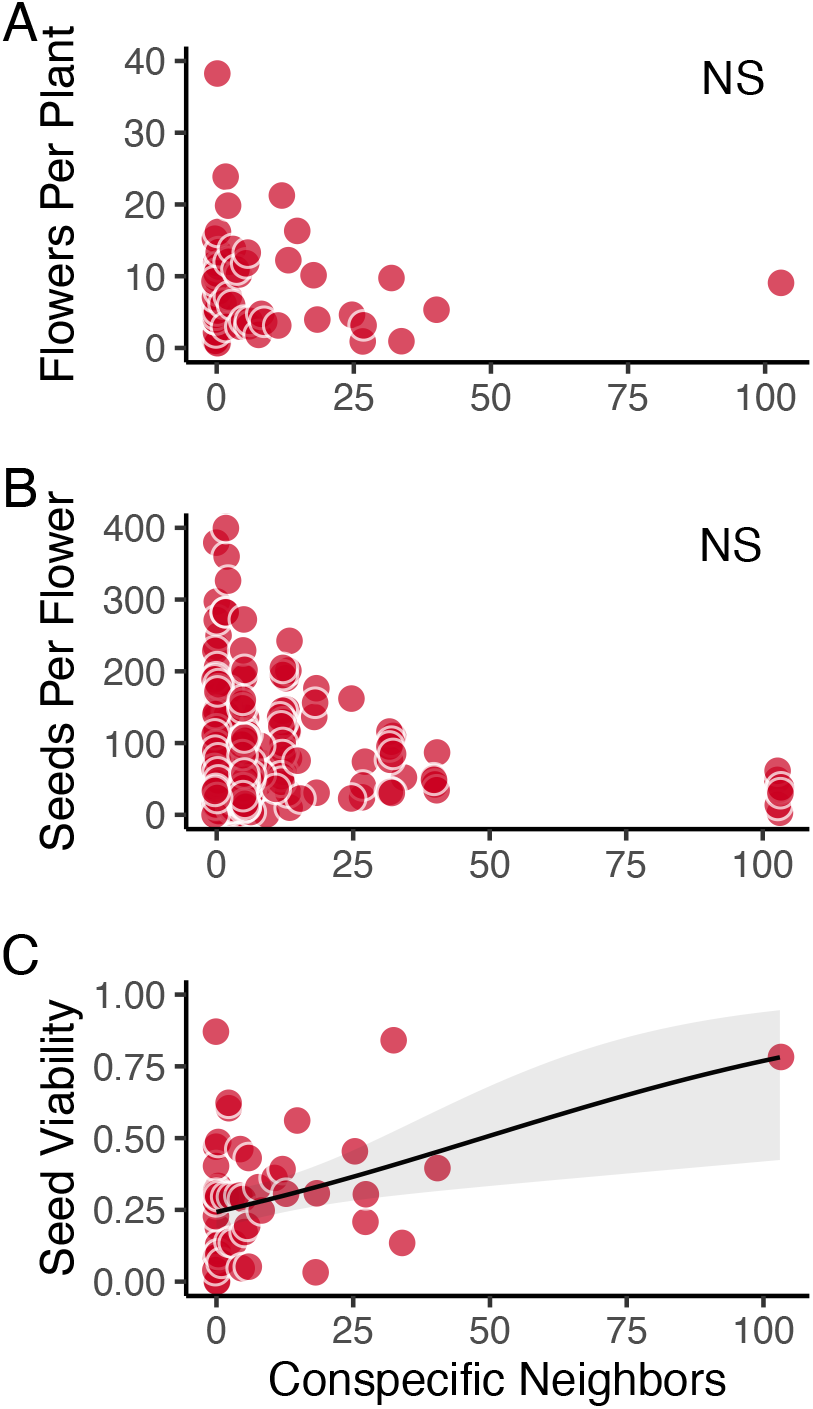
Components of fitness from the 2018 *M. guttatus* immigration experiment. Fitness components: flowers per plant (A), total seeds per flower (B), and seed viability (C). Each red circle represents a single immigrant focal *M. guttatus* transplant in a wash dominated by *M. nudatus*. Conspecific neighbor abundance was significantly associated with focal plant hybridization rate.

Flowers per plant and total seeds per flower for focal immigrant *M. guttatus* transplants were not significantly associated with the number of flowering conspecific neighbor transplants (Figure 3A, 3B; Wald type II Chi-square tests: flowers per plant χ2 = 0.30, df = 1, *p* = 0.58; seeds per flower χ2 = 0.52, df = 1, *p* = 0.47).

One focal immigrant *M. guttatus* transplant had over twice as many surviving neighbors as any other focal transplant. To examine whether including this individual transplant influenced the relationship between fitness components and the number of conspecific neighbors, we reanalyzed the data in two ways: by log transforming the number of neighbors and removing the outlier (Online Supplemental Material: Figures S1 and S2). Neither of these changes qualitatively changed the analysis presented with the raw data.

#### Frequency-manipulation reciprocal transplant

*Fitness components in wash.* For immigrant *M. guttatus* transplants in the wash, seed viability was significantly associated with the frequency treatment (Fig 4E, Wald type II Chisquare test: χ^2^ = 10.602, df = 2, *p* = 0.005). In contrast, flowers per plant (Fig 4A, Wald type II Chi-square test: χ^2^ = 1.75, df = 2, *p* = 0.42) and seeds per flower (Fig 4C, Wald type II Chisquare test: χ^2^ = 0.44, df = 2, *p* = 0.80) were not associated with the frequency treatment. Seed viability in immigrant *M. guttatus* transplants increased from 0.14 to 0.48 between the 5% (1G:18N) and 50% (9G:10N) immigrant frequency treatments (Tukey Post-hoc contrast: *p* = 0.006, Table S1).

**Figure 4.**
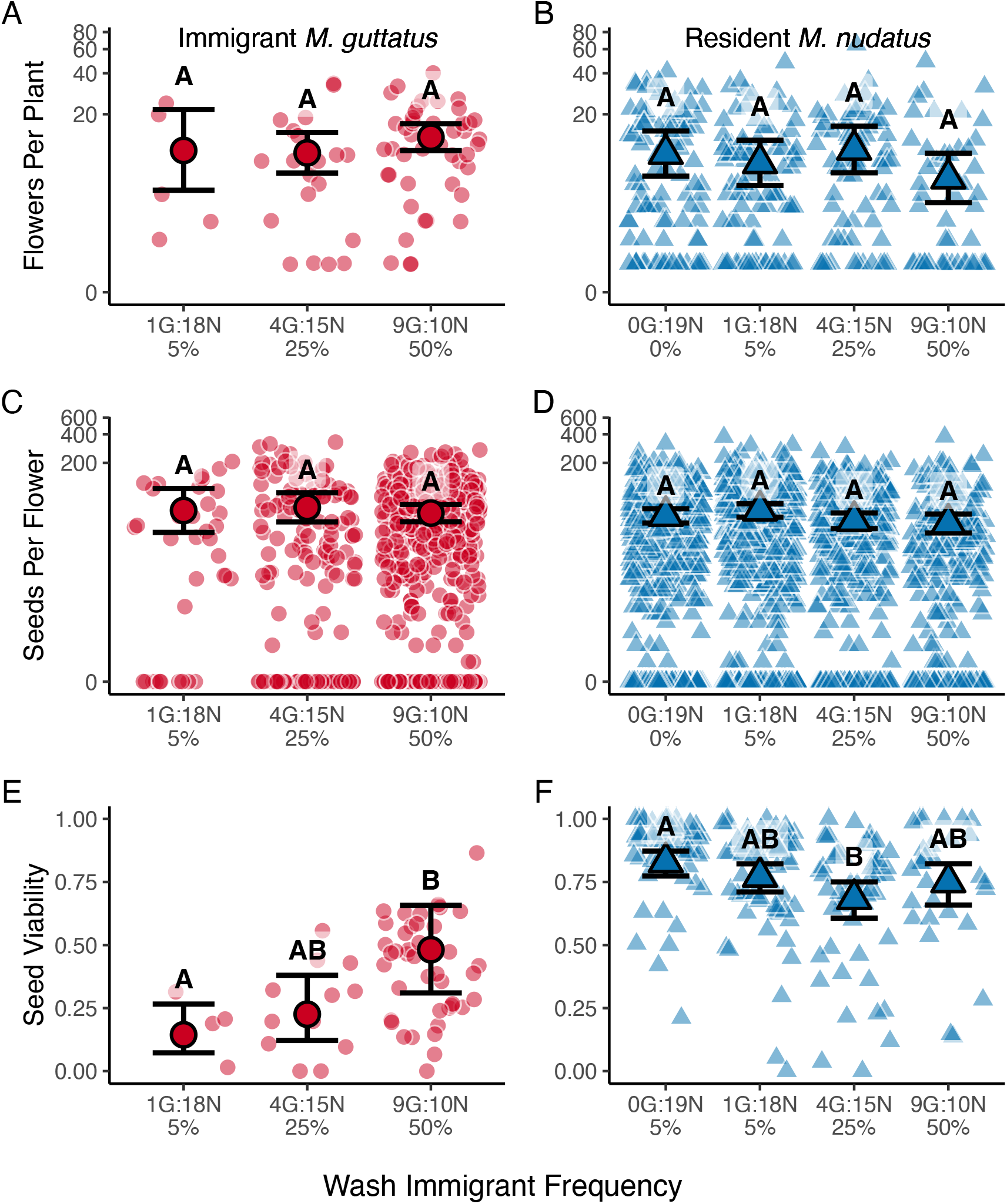
Predicted marginal means and 95% confidence intervals for components of fitness in the wash transplant site from the 2019 reciprocal transplant experiment. Models: Fitness component = immigrant frequency + (1ļBlock). Fitness components: flowers per plant (A, B), total seeds per flower (C, D), and seed viability (E, F). Immigrant *M. guttatus* = red circles (A, C, E), Resident *M. nudatus* = blue triangles (B, D, F). Letters indicate groupings from Tukey post-hoc contrasts.

For resident *M. nudatus* transplants in the wash, seed viability was significantly associated with the frequency treatment (Fig 4F, Wald type II Chi-square test: χ^2^ = 11.40, df = 3, *p* = 0.01), but flowers per plant (Fig 4B, Wald type II Chi-square test: χ^2^ = 3.18, df = 3, *p* =0.37) and seeds per flower were not (Fig 4D, Wald type II Chi-square test: χ^2^ = 6.47, df = 3, *p* = 0.09). Seed viability in resident *M. nudatus* transplants decreased from 0.83 to 0.68 between the 0% (1G:18N) and 25% (9G:10N) immigrant frequency treatments (Tukey Post-hoc contrast: *p* = 0.005, Table S2).

##### Fitness components in seep

For immigrant *M. nudatus* transplants in the seep, flowers per plant (Fig 5A, Wald type II Chi-square test: χ^2^ =11.74, df = 2, *p* = 0.003) and seeds per flower (Fig 5C, Wald type II Chi-square test: χ^2^ = 12.21, df = 2, *p* = 0.002) were significantly associated with frequency treatment, but seed viability was not (Fig 5E, Wald type II Chi-square test: χ^2^ = 1.52, df = 2, *p* = 0.47). Flowers per plant in immigrant *M. nudatus* transplants decreased from ~6 to ~2 flowers per plant between the 25% (4N:15G) and 50% (9N: 10G) immigrant frequency treatments (Tukey post-hoc contrast: *p* = 0.005, Table S3). This decline resulted in a lower proportion of *M. nudatus* flowers than expected in the highest immigrant frequency treatment (Proportion immigrant *M. nudatus* flowers by immigrant frequency treatment: 5% treatment (1N:18G) = 0.06, 25% (4N:15G) treatment = 0.33, 50% (9N:10G) = 0.29). Seeds per flower in immigrant *M. nudatus* transplants increased from 27 to 64 seeds per flower between the 5% (1N:18G) and 25% (4N:15G) immigrant frequency treatments and decreased from 64 to 29 between the 25% (4N:15G) and 50% (9N:10G) immigrant frequency treatments (Tukey post-hoc contrasts: *p* < 0.05, Table S3).

**Figure 5.**
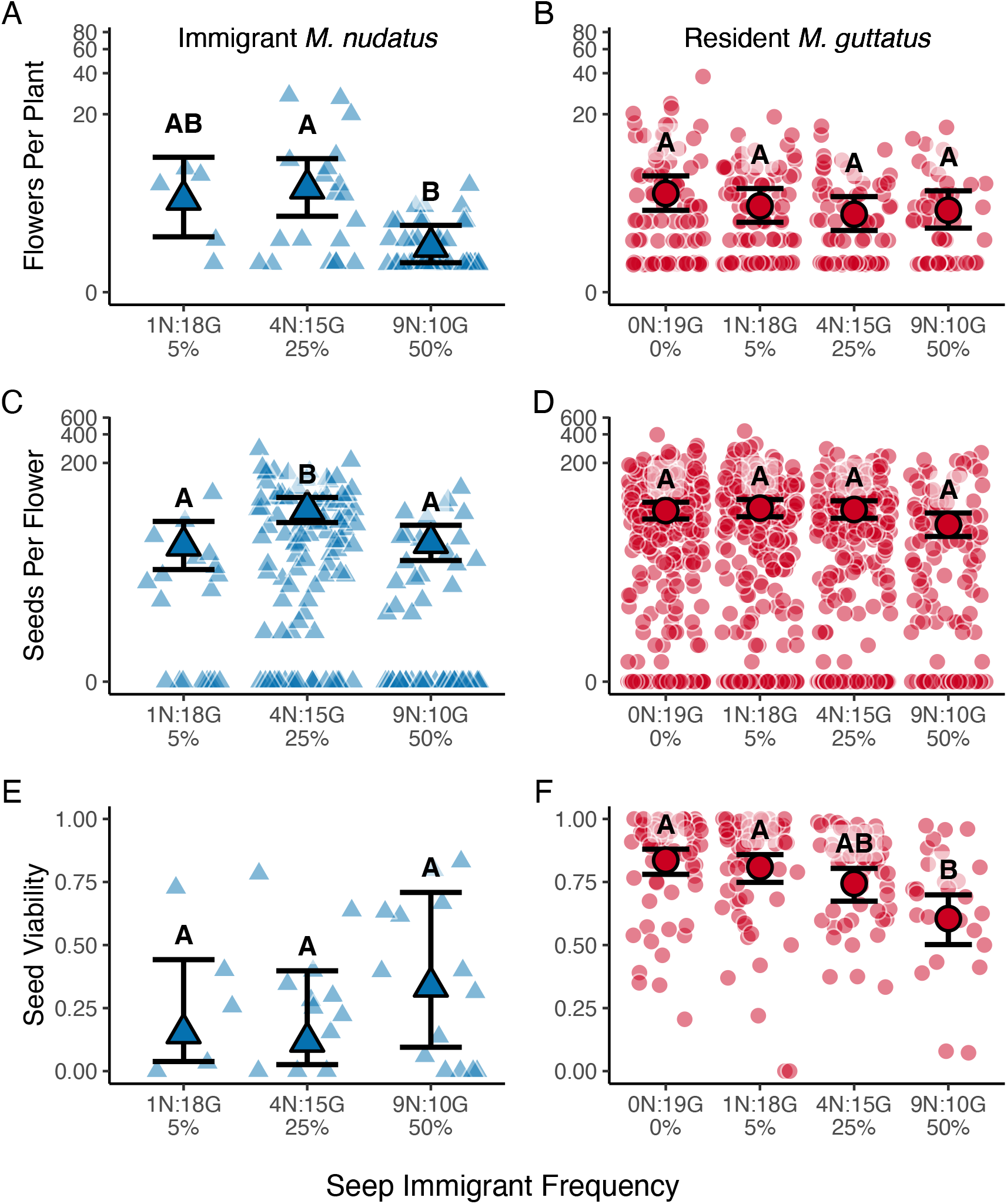
Predicted marginal means and 95% confidence intervals for components of fitness in the seep transplant site from the 2019 reciprocal transplant experiment. Models: Fitness component= immigrant frequency + (1ļBlock). Fitness components: flowers per plant (A, B), total seeds per flower (C, D), and seed viability (E, F). Immigrant *M. nudatus* = blue triangles (A, C, E), resident *M. guttatus* = red circles (B, D, F). Letters indicate groupings from Tukey post-hoc contrasts.

For resident *M. guttatus* transplants in the seep, seed viability was significantly associated with frequency treatment (Fig 5F, Wald type II Chi-square test: χ^2^ = 21.56, df = 3,*p* < 0.001), but flowers per plant (Fig 5B, Wald type II Chi-square test: χ^2^ = 3.14, df = 3, *p* = 0.37) and seeds per flower were not (Fig 5D, Wald type II Chi-square test: χ^2^ = 5.80, df = 3, *p* = 0.122). Seed viability decreased from 0.84 to 0.60 between the 0% (0N:19G) and 50% (9N:10G) immigrant frequency treatments (Tukey post-hoc contrast: *p* < 0.001, Table S4) and decreased from 0.81 to 0.60 between the 5% (1N:18G) and 50% (9N:10G) immigrant frequency treatments (Tukey post-hoc contrast: *p* = 0.001, Table S4)

##### Lifetime fecundity in wash

Viable seeds per plant in the wash was significantly associated with immigrant frequency treatment, species, and an immigrant frequency by species interaction (Fig 6A, Wald type III Chi-square test: frequency χ^2^ = 6.83, df = 2, *p* = 0.03; species χ^2^ = 12.14, df = 1, *p* < 0.001; frequency × species χ^2^ = 14.12, df = 2, *p* < 0.001). Immigrant*M. guttatus* transplants produced one-sixth as many viable seeds per plant as resident *M. nudatus* transplants in the 5% (1N:18G) immigrant frequency treatment (60 vs 347; Tukey post-hoc contrast *p* = 0.008, Table S5).

**Figure 6.**
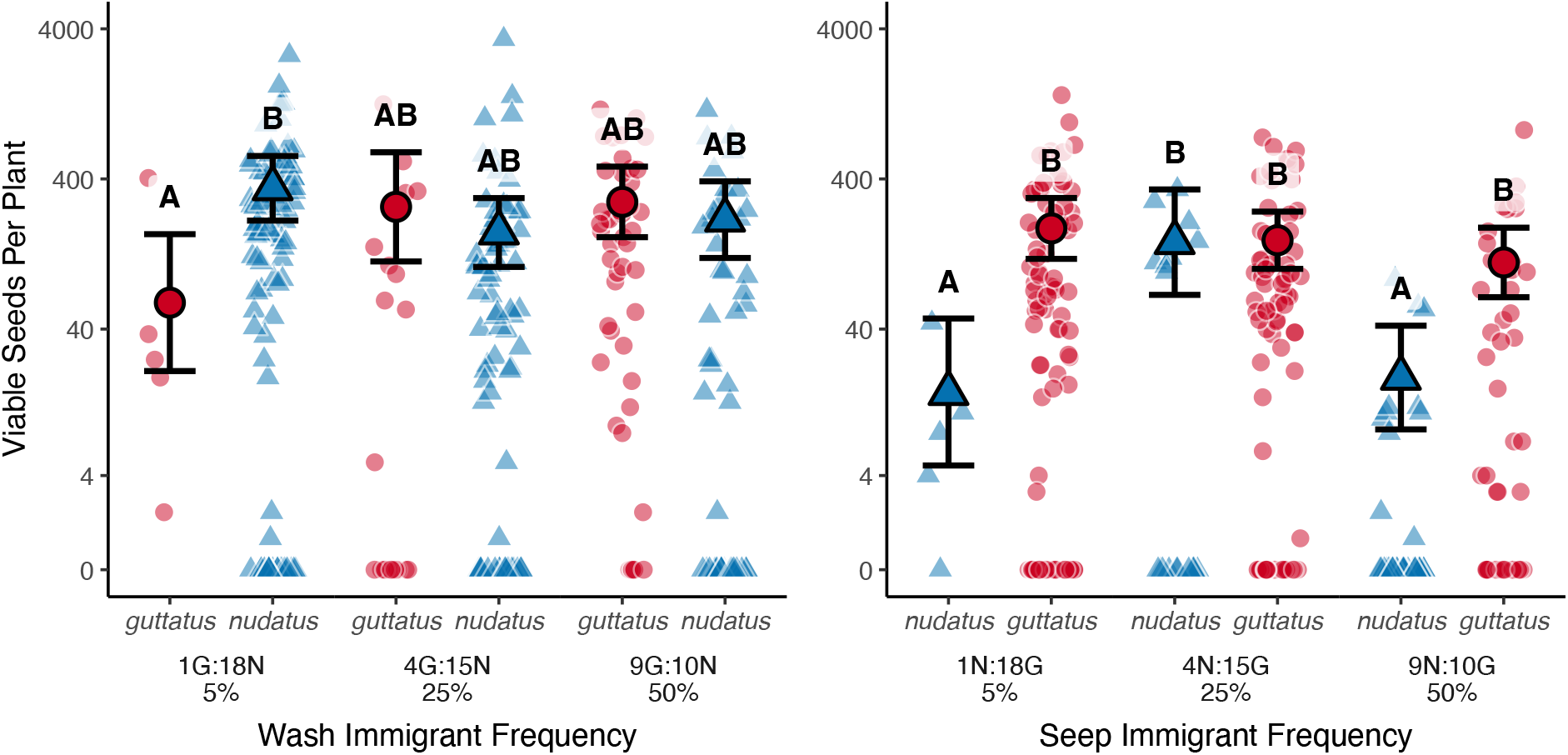
Predicted marginal means and 95% confidence intervals for lifetime fecundity in a wash (A) and seep (B) in the 2019 reciprocal transplant experiment. *M. nudatus* = blue triangles, *M. guttatus* = red circles. Marginal means were predicted from models using data from plots where both species were present at each site. Models: Lifetime fecundity = Species + Immigrant frequency + Species × Immigrant Frequency + (1ļBlock). Letters indicate significant groupings from Tukey post-hoc tests.

##### Lifetime fecundity in seep

Viable seeds per plant in the seep was significantly associated with species and an immigrant frequency treatment by species interaction (Fig 6B, Wald type III Chi-square test: frequency χ^2^ = 2.14, df = 2, p = 0.34; species χ^2^ = 18.85, df = 1,*p* < 0.001; frequency × species χ^2^ = 15.21, df = 2,*p* < 0.001). Immigrant*M. nudatus* transplants produced one-twelfth as many viable seeds per plant as resident *M. guttatus* transplants in the 5% (1N:18G) immigrant frequency treatment (15 vs 187, Tukey post-hoc contrast *p* < 0.001) and one-sixth as many in the 50% (9N:10G) treatment (19 vs 111, Tukey post-hoc contrast *p* = 0.001, Table S6).

#### Experimental Transplant Summary

Seed viability was positively associated with immigrant frequency for immigrant *M. guttatus* transplants, but not immigrant *M. nudatus* transplants, consistent with the findings from the observational transect. Resident transplants of both species also suffered from decreased seed viability as immigrant transplants increased in frequency. Unexpectedly, fitness components declined for immigrant *M. nudatus* transplants when they were most common, resulting in immigrant disadvantage in the highest frequency treatment. Immigrant transplants had lower lifetime fecundity, measured as viable seeds per plant, than resident transplants when immigrants were rarest. For immigrant *M. guttatus*, this rarity disadvantage could not be explained by fitness component differences prior to hybridization. In contrast, rarity disadvantage for immigrant *M. nudatus* was observed for fitness components prior to hybridization.

## Discussion

In this study, we found that frequency-dependent hybridization likely contributes to habitat segregation between two closely related monkeyflower species. Each species suffered from hybridization when in close contact with heterospecifics in both field experiments (Figures 3C, 4E, and 5E), as well as in natural populations (Figure 2). Immigrants of both species had lower lifetime fecundity than residents when rare (Figure 6). However, this minority disadvantage was caused by different factors for each species. Partitioning of the fitness components demonstrated that rarity disadvantage for *M. guttatus* immigrants was most consistent with reproductive interference (Figure 4), whereas aboveground resource and/or pollinator competition together with reproductive interference all likely contributed to rarity disadvantage for *M. nudatus* immigrants (Figure 5). *M. nudatus* immigrants also suffered fitness reductions consistent with intra-specific competition when they were at their highest frequency (Figure 5A). This negative frequency dependence is predicted to facilitate immigrant invasion and break down spatial segregation (Grainger et al. 2019). However, since it was identified only for *M. nudatus,* it is unlikely to lead to breakdown. Our previous studies in this system identified flooding and inbreeding depression as major factors limiting the long-term establishment of *M. nudatus* in habitats occupied by *M. guttatus* (Toll and Willis 2018, Toll et al. 2021). The strong rarity disadvantage for *M. guttatus* immigrants is further evidence that multiple asymmetric processes contribute to the maintenance of strong spatial segregation between these closely related species.

In foundational studies on intertidal algae and barnacles, stress tolerance and competition differed between species, determining zonation patterns (Baker 1909, Connell 1961). The factors limiting each species are mechanistically different in our study, but the asymmetric nature of the limitation is fundamentally similar to the mechanisms of zonation found in those classic studies. For example, we found that a strong barrier limiting *M. nudatus* persistence in wetter sites, occupied by *M. guttatus*, was its inability to survive seasonal flooding (Toll and Willis 2018). Persistent flooding occurs during the winter rainy season in the seeps and meadows inhabited by *M. guttatus*. In contrast, flooding is uncommon in quickly draining washes and rocky outcrops inhabited by *M. nudatus*. An additional asymmetrical barrier limiting immigrant persistence in this system is inbreeding depression (Toll et al. 2021). Since immigrants have fewer opportunities to mate with conspecifics, they likely produce most viable seeds through selfing. While recurrent selfing reduces fitness for both species, the reduction is greater for *M. nudatus* especially in foreign habitats. Selfing reduces immigrant *M. nudatus’* fitness to zero in fewer generations than it does for immigrant *M. guttatus* (Toll et al. 2021). Combined, these prior studies provided a strong mechanistic rationale for why *M. nudatus* does not invade *M. guttatus* streams. However, why *M. guttatus* does not invade *M. nudatus* washes was still largely unexplained, and the experiments presented in this study suggest frequency-dependent hybridization is a contributing factor.

### Frequency dependence of hybridization

This study identifies frequency-dependent hybridization as a strong barrier limiting the persistence of *M. guttatus* in habitats occupied by *M. nudatus* (Figure 4E). This result is similar to that of reciprocal transplant experiments of *Limnanthus* species and *Gilia* subspecies, where frequency-dependent reductions in fecundity for one species (or subspecies) but not the other was attributed to asymmetric impacts of heterospecific pollen transfer on seed set (Nagy 1997, Runquist 2012, Runquist and Stanton 2012). In contrast, Christie and Strauss (2020) observed symmetric declines in seed viability of reciprocally transplanted *Streptanthus* species, even though experimental crosses reduced seed set asymmetrically (Christie and Strauss 2019). Since heterospecific pollen transfer between *M. nudatus* and *M. guttatus* reduces viable seed set symmetrically (Oneal et al. 2016), asymmetry of the frequency-dependence of seed viability is surprising. However, this asymmetry can be explained when we consider how fecundity components prior to hybridization changed across immigrant frequency treatments for *M. nudatus* immigrants. *M. nudatus* immigrants produced the fewest flowers per plant in the highest immigrant frequency treatment, resulting in a lower proportion of *M. nudatus* gametes than suggested by the plant frequency treatment (~50% plants vs ~29% flowers). *M. nudatus* has fewer ovules and pollen grains per flower than *M. guttatus* (Oneal et al. 2016, Ritland and Ritland 1989). This difference in pollen production further amplified the difference in effective gamete frequency than would be expected solely by the manipulation of the frequency of *M. nudatus*. Thus, while other studies were able to predict asymmetries in frequency-dependence based on lab crossing data (Runquist and Stanton 2012, Nagy 1997), the asymmetry we observe is driven by ecological interactions (the reduction in flower number with increasing conspecific frequency) and evolved life history differences (pollen production).

While naturally occurring *M. nudatus* also hybridizes with *M. guttatus,* seed viability was also not associated with conspecific fruit frequency in the observational transect (Figure 2). Our result differs from that of Gardner and MacNair (2000), who found a significant negative correlation between fruit frequency and seed viability in *M. nudatus* co-occuring with *M. guttatus*. Our studies occurred in the same region but in qualitatively different sites. Gardner and MacNair (2000) only sampled *M. nudatus* plants with heterospecifics in close proximity, equivalent to the middle portion of our transect where both species co-occurred but did not extend the transect into pure populations. Compared to *M. guttatus*, seed viability in the transition zone for *M. nudatus* was more variable (Figure 2), but since we did not record phenology or pollinator behavior over time, the factors contributing to this pattern remain open questions. While the onset of flowering differs between these species in some sites or experimental conditions (Selby et al. 2014, Sianta and Kay 2021, Wu et al. 2010), these species have highly overlapping flowering periods (Gardner and MacNair 2000, personal observation, 2013-2019). Further, the relative order and even whether the onset of flowering differs between species is site and daylength dependent (Friedman and Willis 2013, Toll 2017, K. Toll and J. H. Willis unpublished data).

Our observations from the transect study suggest that fitness costs of hybridization are highly localized and decay within meters from the nearest heterospecific (Figure 2), which makes sense given the narrow boundaries that we see in most populations. In the reciprocal transplant experiment, resident transplants of both species suffered decreased seed viability when immigrant transplants were at high frequencies (Figure 4F, 5F), despite the high abundance of native conspecifics for resident transplants at each site. These results suggest that low pollinator constancy leads to high rates of interspecific pollen transfer between species at the local scale, and that most pollination is occurring at a very small scale. These localized and frequencydependent fitness costs to residents are consistent with the observed stability of spatial segregation patterns. Immigration rates would have to be extremely high, and immigrants would have to be widespread to negatively affect resident populations.

### Competition may contribute asymmetrically to habitat segregation

Competition could promote or limit coexistence depending on the degree of niche overlap between species and whether any species is competitively dominant (Chesson 2000). In the absence of strong intraspecific competition, if niche overlap between the species is high, immigrant fitness components would be positively associated with immigrant frequency, potentially resulting in competitive exclusion of the immigrant species (Gause 1934, Hardin 1961). It is possible that one species could always be competitively dominant over the other, or dominance could be context dependent and contribute asymmetrically to habitat segregation (Bertness 1991, Connell 1961, Grace and Wetzel 1981). However, since intraspecific competition is usually stronger than interspecific competition (Adler et al. 2018), rare immigrants may instead experience competitive release in habitats lacking conspecifics. If niche overlap between species is low, immigrant fitness components would be negatively associated with immigrant frequency, requiring greater fitness costs of hybridization to counterbalance the negative frequency dependence that would promote local coexistence and erode spatial segregation (Adler et al. 2007, Adler et al. 2018).

The *M. guttatus/M. nudatus* study system presented a unique opportunity to separate effects of hybridization on fitness from effects of competition and abiotic factors. While the combined effects of reproductive and ecological niche overlap could amplify positive frequency dependence and hasten exclusion (Schreiber et al. 2019), we instead observed declines in flowers per plant and seeds per flower for *M. nudatus* immigrants in their highest frequency treatment. The decline at high densities suggests that some form of competition was more important than release from reproductive interference for *M. nudatus* immigrants at high immigrant frequencies. As the transplants occurred in plastic conetainers, pre-empting below-ground competition, above-ground competition for space, light, and/or pollinator visits may have suppressed flower and seed production in these potted immigrants (Figure 1D). Due to the experimental design of our frequency arrays (Figure 1C), we were unable to estimate the strength of intra-vs interspecific competition (Inouye 2001).

Immigrant *M. nudatus* transplants produced the greatest number of seeds per flower in the intermediate immigrant frequency treatment (Figure 5C). We do not have a clear interpretation for this phenomenon. However, we hypothesize that pollinator behavior might have driven this U-shaped pattern in fecundity. Benadi and Pauw (2018) observed that South African Fynbos species received the highest visitation rates at intermediate flower abundances, relative to rare and common species. Future studies on pollinator visitation rates are needed to test this hypothesis. *M. guttatus* and *M. nudatus* are visited by many overlapping generalist bee species, but *M. nudatus* visitors tend to be small-bodied while *M. guttatus* visitors are small to large-bodied (Gardner and MacNair 2000, Koski et al. 2015, K. Toll and D. B. Lowry, unpublished data).

In greenhouse studies, Brigham (2003) failed to find deleterious effects of interspecific competition to either *M. guttatus* or *M. nudatus*. In the field, the effects of intra-specific competition for *M. nudatus* varied by year, site, and the timing of germination, resulting in conspecific neighbor removals having positive, negative, or no effect on *M. nudatus* survival (Brigham 2001). In our study, we did not observe reductions in *M. guttatus* fecundity prior to hybridization (i.e., in flower or fruit set). These findings accord well with expected competitive ability of *M. guttatus,* which occur in streams that are full of both conspecifics as well as a high density of many other plant species. The rocky serpentine washes inhabited by *M. nudatus* are mostly sparsely vegetated, with only occasional areas of high density of monkeyflowers and very few other small forbs. This is consistent with the broader pattern of serpentine endemics occurring in barer (low competition) habitats compared to closely related species (Sianta and Kay 2019). Therefore, it is perhaps not surprising that in our arrays, which intentionally created a density somewhat intermediate and appropriate for both habitats, *M. nudatus* would suffer from competition.

### Reciprocal transplant experiments and the asymmetry of barriers

Reciprocal transplant experiments provide strong evidence of local adaptation, whether between populations, ecotypes, or closely related species (Clausen et al. 1940, Kawecki and Ebert 2004, Hereford 2009). In annual plants, the response variable in these studies is usually seed set. Disentangling the relative effects of interspecific mating from environmental differences between habitats can be achieved by varying the frequencies of each species reciprocally in native and foreign sites. This allows comparisons across frequencies of fitness components that are not affected by interspecific mating interactions (e.g., survival, flower number) to those that could be (e.g., seed abortion, hybrid seed production, Nagy 1997, Runquist and Stanton 2013, Christie and Strauss 2020).

In reciprocal transplants, the fitness of local residents and foreign immigrants are compared at native and foreign sites and changes in the rank of fitness, where local residents outperform foreign immigrants at each site, is taken as evidence of adaptation to local conditions (Kawecki and Ebert 2004). Beyond differential habitat-based selection, transplanting individuals of closely related species reciprocally reduces the distance between heterospecific individuals. This in turn increases the chances of reproductive interference. While reciprocal transplant experiments are often used to explain habitat segregation between closely related species (e.g., Yost et al. 2012, Peterson et al. 2013, Chen and Schemske 2015, Ferris and Willis 2018, DiVittorio et al. 2020), reproductive interactions are rarely considered as important drivers of these patterns (but see Toll and Willis 2018, Christie and Strauss 2020). Since heterospecific pollen transfer can reduce seed set in a pattern indistinguishable from divergent adaptation, deleterious effects of interspecific mating on fitness need to be accounted for in assessments of local adaptation.

As demonstrated here, both divergent habitat adaptation (i.e., differences in flooding tolerance) and reproductive interference contribute to the spatial separation of the two monkeyflower species. The underlying mechanisms of resident species advantage in reciprocal transplants commonly operate asymmetrically (Popovic and Lowry 2020), and thus a pattern consistent with divergent adaptation may arise as an emergent property of several asymmetric barriers to foreign habitat persistence. Therefore, interpretations of a single fitness metrics (i.e., flower production, seed set) across species are difficult to interpret without knowledge of frequency- and density-dependent processes and the underlying mechanisms. In a previous study, *M. guttatus* and *M. nudatus* were transplanted reciprocally at equal frequencies (Toll and Willis 2018) and thus, could not test for minority disadvantage as we did here.

## Conclusion

Patterns of spatial segregation are ubiquitous across animal and plant taxa and diverse environments (Wisheu 1998). In closely related species, negative reproductive interactions can contribute to these patterns if they occur in a frequency-dependent manner and are stronger than intraspecific competition (Schreiber et al 2019, Weber and Strauss 2016). Using both experimental and observational approaches, we found that hybridization operated in a frequencydependent manner for one species of monkeyflower when invading habitats dominated by its congener but was not frequency-dependent when the other species invaded. Despite this asymmetry, hybridization contributed to rare immigrant disadvantage for both species. Overall, this result, combined with other results from this system (Toll and Willis 2018, Toll et al. 2021), suggests that spatial segregation is an emergent property of many asymmetric barriers operating in complementary and contrasting ways. Therefore, it seems likely that evidence of divergent adaptation from reciprocal transplant experiments of closely related species have the potential to be driven by reproductive interference, in addition to the more commonly invoked differences in stress tolerance and competitive ability. Parsing out the specific effects of each mechanism across many systems will be necessary for a fuller understanding of why closely related species segregate into different habitats.

## Acknowledgments

The authors wish to thank K. Christie, J.M. Coughlan, and E.F. LoPresti whose comments helped improve and clarify this manuscript. This work was performed at the University of California Natural Reserve System McLaughlin Natural Reserve, Reserve DOI: 10.21973/N3W08D. We thank C. Koehler and P. Aigner for facilitating field work. This work was supported by Michigan State University and a grant from the National Science Foundation (IOS-1855927).

## Statement of Authorship

KT and DBL conceived of the study. KT designed and performed the field experiments and collected and analyzed the data. KT and DBL wrote and edited the manuscript.

## Data and Code Accessibility

Data and R code are accessible through Dryad (doi: 10.5061/dryad.d51c5b04g; Toll and Lowry 2022) and Zenodo (doi: 10.5281/zenodo.5829342).

## Notes

https://doi.org/10.5061/dryad.d51c5b04g

### Competing Interest Statement

The authors have declared no competing interest.

