## SupplementalMaterials for "Frequency-dependent hybridization contributes to habitat segregation in monkeyflowers"

Katherine Toll[1,*]

David B. Lowry[1,2]

1. Department of Plant Biology, Michigan State University, East Lansing, Michigan 48824
2. Program in Ecology, Evolution, and Behavior, Michigan State University, East Lansing, Michigan 48824

*Keywords*: frequency-dependence, hybridization, reproductive interference, habitat segregation, *Mimulus*, monkeyflower

Table S1. Tukey post-hoc contrasts for fitness component models of immigrant *M. guttatus* in the wash in 2019. Fitness component = immigrant frequency treatment + (1|block). *P*-values < 0.05 in bold.

| Fitness component | contrast | estimate | SE | *df* | *t*-ratio | *p*-value |
|---|---|---|---|---|---|---|
| Flowers per plant | 5% vs 25% | 0.0426 | 0.397 | 65 | 0.107 | 0.9937 |
| | 5% vs 50% | -0.2256 | 0.373 | 65 | -0.604 | 0.8183 |
| | 25% vs 50% | -0.2682 | 0.213 | 65 | -1.261 | 0.4223 |
| Seeds per flower | 5% vs 25% | -0.0761 | 0.325 | 863 | -0.234 | 0.9703 |
| | 5% vs 50% | 0.0629 | 0.291 | 863 | 0.216 | 0.9747 |
| | 25% vs 50% | 0.139 | 0.212 | 863 | 0.656 | 0.7889 |
| Seed viability | 5% vs 25% | -0.55 | 0.543 | 66 | -1.013 | 0.5713 |
| | 5% vs 50% | -1.71 | 0.539 | 66 | -3.171 | **0.0064** |
| | 25% vs 50% | -1.16 | 0.53 | 66 | -2.186 | 0.0811 |

Table S2. Tukey post-hoc contrasts for fitness component models of resident *M. nudatus* in the wash in 2019. Fecundity component = immigrant frequency treatment + (1|block). *P*-values < 0.05 in bold.

| Fitness component | contrast | estimate | SE | *df* | *t*-ratio | *p*-value |
|---|---|---|---|---|---|---|
| Flowers per plant | 0% vs 5% | 0.1602 | 0.282 | 304 | 0.568 | 0.9415 |
| | 0% vs 25% | -0.0702 | 0.285 | 304 | -0.247 | 0.9947 |
| | 0% vs 50% | 0.4265 | 0.298 | 304 | 1.431 | 0.4807 |
| | 5% vs 25% | -0.2304 | 0.286 | 304 | -0.806 | 0.8516 |
| | 5% vs 50% | 0.2663 | 0.299 | 304 | 0.890 | 0.8099 |
| | 25% vs 50% | 0.4967 | 0.302 | 304 | 1.646 | 0.3543 |
| Seeds per flower | 0% vs 5% | -0.132 | 0.121 | 3100 | -1.087 | 0.6978 |
| | 0% vs 25% | 0.12 | 0.13 | 3100 | 0.919 | 0.7948 |
| | 0% vs 50% | 0.186 | 0.146 | 3100 | 1.273 | 0.5804 |
| | 5% vs 25% | 0.251 | 0.128 | 3100 | 1.966 | 0.2012 |
| | 5% vs 50% | 0.317 | 0.143 | 3100 | 2.221 | 0.1177 |
| | 25% vs 50% | 0.066 | 0.152 | 3100 | 0.435 | 0.9724 |
| Seed viability | 0% vs 5% | 0.362 | 0.234 | 304 | 1.551 | 0.4083 |
| | 0% vs 25% | 0.81 | 0.242 | 304 | 3.349 | **0.005** |
| | 0% vs 50% | 0.482 | 0.282 | 304 | 1.708 | 0.3214 |
| | 5% vs 25% | 0.448 | 0.231 | 304 | 1.938 | 0.2144 |
| | 5% vs 50% | 0.12 | 0.273 | 304 | 0.438 | 0.9718 |
| | 25% vs 50% | -0.328 | 0.279 | 304 | -1.176 | 0.6427 |

Table S3. Tukey post-hoc contrasts for fitness components models of immigrant *M. nudatus* in the seep in 2019. Fecundity component = immigrant frequency treatment + (1|block). *P*-values < 0.05 in bold.

| Fitness component | contrast | estimate | SE | *df* | *t*-ratio | *p*-value |
|---|---|---|---|---|---|---|
| Flowers per plant | 5% vs 25% | -0.205 | 0.461 | 65 | -0.444 | 0.8972 |
| | 5% vs 50% | 0.989 | 0.456 | 65 | 2.169 | 0.0843 |
| | 25% vs 50% | 1.194 | 0.364 | 65 | 3.283 | **0.0047** |
| Seeds per flower | 5% vs 25% | -0.8619 | 0.336 | 233 | -2.562 | **0.0296** |
| | 5% vs 50% | -0.0639 | 0.371 | 233 | -0.172 | 0.9838 |
| | 25% vs 50% | 0.798 | 0.269 | 233 | 2.969 | **0.0092** |
| Seed viability | 5% vs 25% | 0.289 | 1.11 | 66 | 0.262 | 0.9629 |
| | 5% vs 50% | -1.048 | 1.11 | 66 | -0.945 | 0.6138 |
| | 25% vs 50% | -1.337 | 1.15 | 66 | -1.166 | 0.4776 |

Table S4. Tukey post-hoc contrasts for fitness component models of resident *M. guttatus* in the seep in 2019. Fitness component = immigrant frequency treatment + (1|block). *P*-values < 0.05 in bold.

| Fitness component | contrast | estimate | SE | *df* | *t*-ratio | *p*-value |
|---|---|---|---|---|---|---|
| Flowers per plant | 0% vs 5% | 0.2292 | 0.228 | 304 | 1.007 | 0.7456 |
| | 0% vs 25% | 0.3883 | 0.232 | 304 | 1.671 | 0.341 |
| | 0% vs 50% | 0.3107 | 0.243 | 304 | 1.281 | 0.576 |
| | 5% vs 25% | 0.1592 | 0.234 | 304 | 0.679 | 0.9051 |
| | 5% vs 50% | 0.0816 | 0.245 | 304 | 0.333 | 0.9872 |
| | 25% vs 50% | -0.0776 | 0.249 | 304 | -0.312 | 0.9895 |
| Seeds per flower | 0% vs 5% | -0.0599 | 0.147 | 1347 | -0.408 | 0.9771 |
| | 0% vs 25% | -0.0261 | 0.149 | 1347 | -0.175 | 0.9981 |
| | 0% vs 50% | 0.3426 | 0.176 | 1347 | 1.948 | 0.2087 |
| | 5% vs 25% | 0.0338 | 0.151 | 1347 | 0.224 | 0.9961 |
| | 5% vs 50% | 0.4024 | 0.179 | 1347 | 2.252 | 0.11 |
| | 25% vs 50% | 0.3686 | 0.18 | 1347 | 2.047 | 0.1713 |
| Seed viability | 0% vs 5% | 0.183 | 0.251 | 304 | 0.728 | 0.886 |
| | 0% vs 25% | 0.562 | 0.248 | 304 | 2.269 | 0.1077 |
| | 0% vs 50% | 1.205 | 0.278 | 304 | 4.328 | **0.0001** |
| | 5% vs 25% | 0.379 | 0.244 | 304 | 1.553 | 0.4073 |
| | 5% vs 50% | 1.022 | 0.276 | 304 | 3.704 | **0.0014** |
| | 25% vs 50% | 0.644 | 0.272 | 304 | 2.364 | 0.0863 |

Table S5. Tukey post hoc contrasts for lifetime fecundity model in the wash transplant site. IFT= immigrant frequency treatment. Lifetime fecundity = immigrant frequency treatment + species + treatment*species + (1|block). *P*-values < 0.05 in bold.

| contrast | estimate | SE | *df* | *t*-ratio | *p*-value |
|---|---|---|---|---|---|
| *M. guttatus* (5% IFT) - *M. nudatus* (5% IFT) | -1.7539 | 0.503 | 276 | -3.484 | **0.0075** |
| *M. guttatus* (5% IFT) - *M. guttatus* (25% IFT) | -1.4696 | 0.685 | 276 | -2.147 | 0.2665 |
| *M. guttatus* (5% IFT) - *M. nudatus* (25% IFT) | -1.0762 | 0.598 | 276 | -1.799 | 0.4682 |
| *M. guttatus* (5% IFT) - *M. guttatus* (50% IFT) | -1.5457 | 0.603 | 276 | -2.563 | 0.11 |
| *M. guttatus* (5% IFT) - *M. nudatus* (50% IFT) | -1.2732 | 0.614 | 276 | -2.073 | 0.3044 |
| *M. nudatus* (5% IFT) - *M. guttatus* (25% IFT) | 0.2843 | 0.496 | 276 | 0.573 | 0.9927 |
| *M. nudatus* (5% IFT) - *M. nudatus* (25% IFT) | 0.6777 | 0.369 | 276 | 1.835 | 0.4453 |
| *M. nudatus* (5% IFT) - *M. guttatus* (50% IFT) | 0.2081 | 0.374 | 276 | 0.556 | 0.9936 |
| *M. nudatus* (5% IFT) - *M. nudatus* (50% IFT) | 0.4807 | 0.392 | 276 | 1.226 | 0.824 |
| *M. guttatus* (25% IFT) - *M. nudatus* (25% IFT) | 0.3933 | 0.396 | 276 | 0.994 | 0.9197 |
| *M. guttatus* (25% IFT) - *M. guttatus* (50% IFT) | -0.0762 | 0.509 | 276 | -0.15 | 1 |
| *M. guttatus* (25% IFT) - *M. nudatus* (50% IFT) | 0.1964 | 0.522 | 276 | 0.376 | 0.999 |
| *M. nudatus* (25% IFT) - *M. guttatus* (50% IFT) | -0.4695 | 0.386 | 276 | -1.218 | 0.828 |
| *M. nudatus* (25% IFT) - *M. nudatus* (50% IFT) | -0.197 | 0.403 | 276 | -0.489 | 0.9965 |
| *M. guttatus* (50% IFT) - *M. nudatus* (50% IFT) | 0.2725 | 0.269 | 276 | 1.015 | 0.9128 |

Table S6. Tukey post hoc contrasts for lifetime fecundity model in the seep transplant site. IFT= immigrant frequency treatment. Lifetime fecundity = immigrant frequency treatment + species + treatment*species + (1|block). *P*-values < 0.05 in bold.

| contrast | estimate | SE | *df* | *t*-ratio | *p*-value |
|---|---|---|---|---|---|
| *M. guttatus* (5% IFT) - *M. nudatus* (5% IFT) | 2.529 | 0.582 | 276 | 4.342 | **0.0003** |
| *M. guttatus* (5% IFT) - *M. guttatus* (25% IFT) | 0.1813 | 0.325 | 276 | 0.558 | 0.9936 |
| *M. guttatus* (5% IFT) - *M. nudatus* (25% IFT) | 0.2107 | 0.47 | 276 | 0.448 | 0.9977 |
| *M. guttatus* (5% IFT) - *M. guttatus* (50% IFT) | 0.5218 | 0.358 | 276 | 1.456 | 0.6924 |
| *M. guttatus* (5% IFT) - *M. nudatus* (50% IFT) | 2.2937 | 0.48 | 276 | 4.775 | **<.0001** |
| *M. nudatus* (5% IFT) - *M. guttatus* (25% IFT) | -2.3478 | 0.627 | 276 | -3.747 | **0.003** |
| *M. nudatus* (5% IFT) - *M. nudatus* (25% IFT) | -2.3183 | 0.715 | 276 | -3.242 | 0.0166 |
| *M. nudatus* (5% IFT) - *M. guttatus* (50% IFT) | -2.0073 | 0.645 | 276 | -3.111 | 0.0249 |
| *M. nudatus* (5% IFT) - *M. nudatus* (50% IFT) | -0.2354 | 0.715 | 276 | -0.329 | 0.9995 |
| *M. guttatus* (25% IFT) - *M. nudatus* (25% IFT) | 0.0295 | 0.399 | 276 | 0.074 | 1 |
| *M. guttatus* (25% IFT) - *M. guttatus* (50% IFT) | 0.3405 | 0.352 | 276 | 0.968 | 0.9277 |
| *M. guttatus* (25% IFT) - *M. nudatus* (50% IFT) | 2.1124 | 0.467 | 276 | 4.519 | **0.0001** |
| *M. nudatus* (25% IFT) - *M. guttatus* (50% IFT) | 0.3111 | 0.493 | 276 | 0.632 | 0.9886 |
| *M. nudatus* (25% IFT) - *M. nudatus* (50% IFT) | 2.083 | 0.584 | 276 | 3.568 | **0.0056** |
| *M. guttatus* (50% IFT) - *M. nudatus* (50% IFT) | 1.7719 | 0.443 | 276 | 3.998 | **0.0011** |

*M. guttatus* immigration experiment: Sensitivity analysis

One focal plant had 2.5 times more neighbors (103) than focal plant with the next highest number of neighbors (40). To examine whether this individual influenced the conclusions of our analysis, we repeated the statistical analysis of the *M. guttatus immigration* experiment in two ways: 1) log transforming (log(Conspecific Neighbors +1)) the number of conspecific neighbors and 2) removing the outlier from the analysis. These analyses do not qualitatively change the results presented in the manuscript.

*Log transformation*. Seed viability increased with increasing log transformed conspecific neighbors (Wald type II Chi-square test: $\chi 2 = 7.95$, df = 1, $p = 0.004$; coefficient estimate = 0.28, SE = 0.099, z-value = 2.82, $p = 0.005$). The model-predicted seed viability for rare immigrants increased from 0.22 for immigrants without log transformed conspecific neighbors, to 0.50 in the plant with 4.6 log transformed neighbors (Figure S1 C).

Flowers per plant and total seeds per flower for focal immigrant *M. guttatus* transplants were not significantly associated with the log transformed number of flowering conspecific neighbor transplants (Figure S1 A, S1 B; Wald type II Chi-square tests: flowers per plant $\chi 2 = 0.03$, df = 1, $p = 0.87$; seeds per flower $\chi 2 = 0.34$, df = 1, $p = 0.56$).

*Removing outlier.* Seed viability increased with increasing log transformed conspecific neighbors (Wald type II Chi-square test: $\chi 2 = 4.28$, df = 1, $p = 0.039$; coefficient estimate = 0.026, SE = 0.013, z-value = 2.07, $p = 0.039$). The model-predicted seed viability for rare immigrants increased from 0.24 for immigrants without conspecific neighbors, to 0.47 in the plant with 40 neighbors (Figure S2 C).

Flowers per plant and total seeds per flower for focal immigrant *M. guttatus* transplants were not significantly associated with the log transformed number of flowering conspecific neighbor transplants (Figure S2 A, S2 B; Wald type II Chi-square tests: flowers per plant $\chi 2$ = 1.57, df = 1, $p$ = 0.21; seeds per flower $\chi 2$ = 0.32, df = 1, $p$ = 0.57).

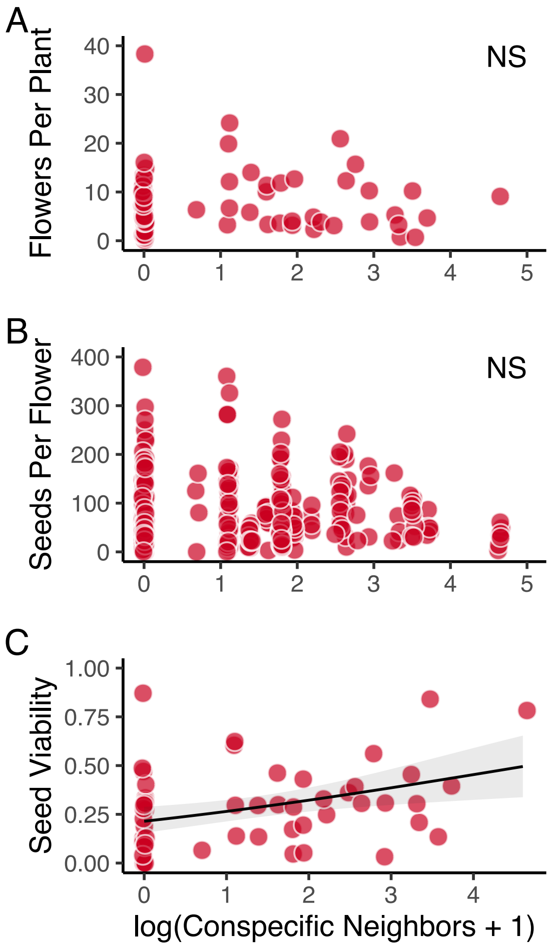


Figure S1. Components of fitness from the 2018 *M. guttatus* immigration experiment, with conspecific neighbors log transformed. Fitness components: flowers per plant (A), total seeds per flower (B), and seed viability (C). Each red circle represents a single immigrant focal *M. guttatus* transplant in a wash dominated by *M. nudatus*. Log transformed conspecific neighbor abundance was significantly associated with focal plant hybridization rate.

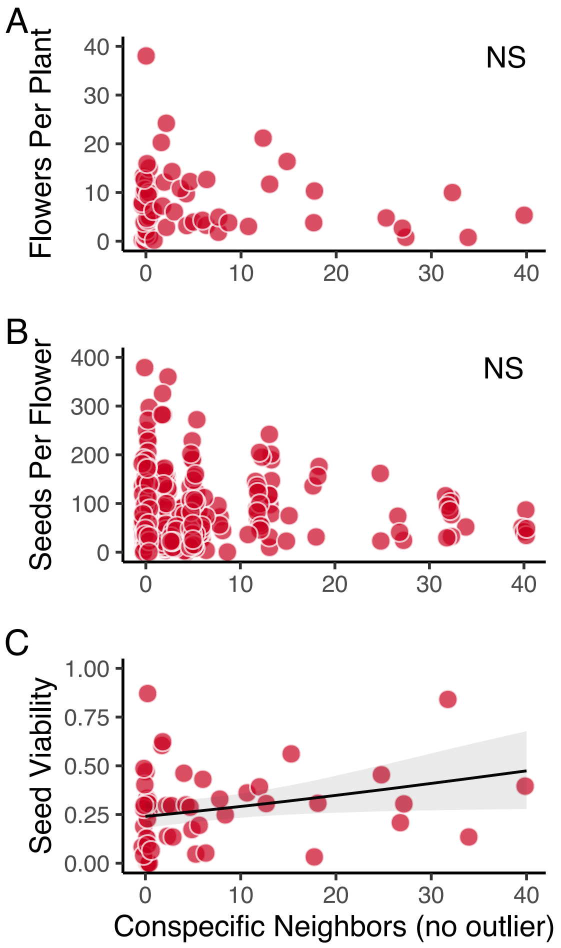


Figure S2. Components of fitness from the 2018 *M. guttatus* immigration experiment, omitting an outlier focal plant with 103 neighbors. Fitness components: flowers per plant (A), total seeds per flower (B), and seed viability (C). Each red circle represents a single immigrant focal *M. guttatus* transplant in a wash dominated by *M. nudatus*. Conspecific neighbor abundance was significantly associated with focal plant hybridization rate
